## Supplemental Materials for "Fast holographic scattering compensation for deep tissue biological imaging"

### I. SPEED AND PHOTON BUDGET COMPARISON

The fast convergence characteristics of DASH allow for finding scattering correction patterns in fewer measurements and using fewer photons compared to other methods. Here we quantitatively investigate the total measurement times and photon budgets required to establish a correction level of 50%. We assume a homogeneous 2D fluorescent layer as the sample, 256 scattered and the same number of correctable modes. Assuming matching mode numbers here allows us to easily determine the signal at optimal correction by simulating an undisturbed focus. The signal flux is assumed to be around 2000 photons per 100 µs dwell time if the scattering is not corrected.

We consider different correction strategies and SLM variants, which differ in their pixel transition times. For F-SHARP, no time is added for mode changing and phase stepping, because changing the plane wave modes is done via galvo mirrors and the number of required phase stepping procedures is negligible (only one step after one full galvo scan). We note that this is a very optimistic assumption, as the inertia of any mechanical scanning device will in practice cause delays, too. For DASH we consider different SLM types with increasing refresh rates $t_{SLM}$: a fast nematic LC-SLM ($t_{SLM} = 3$ ms), a fast ferroelectric LC-SLM ($t_{SLM} = 0.2$ ms), and a fast MEMS mirror with a switching time of 50 µs, for which we assumed an additional pause of 50 µs whenever an updated correction pattern is calculated (regularly after groups of three measurements taken at different phase-steps). For the ferroelectric SLM we further considered the fact that the current technology does not allow for direct updating at a 5 kHz rate. Instead, individual frames consisting of 24 bitplanes are transferred to the SLM at a correspondingly lower rate, which are then displayed in a fast sequence. This means that a feedback based update of the correction pattern can only occur after a set comprising 8 modes (each measured at 3 phase steps = 24 images) has been tested.

Fig. S1 summarizes the results. Each data point is the average result obtained in 5 subsequent simulated measurements on different random scatterers. The left graph visualizes the photon budget required to achieve a correction level of 50%, at which the TPEF fluorescence image of a 2D layer appears at 50% brightness compared to the undisturbed case. The simulation includes Poissonian shot noise, but otherwise assumes an ideal, noise-free detector. Apparently, all DASH implementations require roughly half as many photons than F-SHARP, which is explained by the higher information conveyed per detected photon.

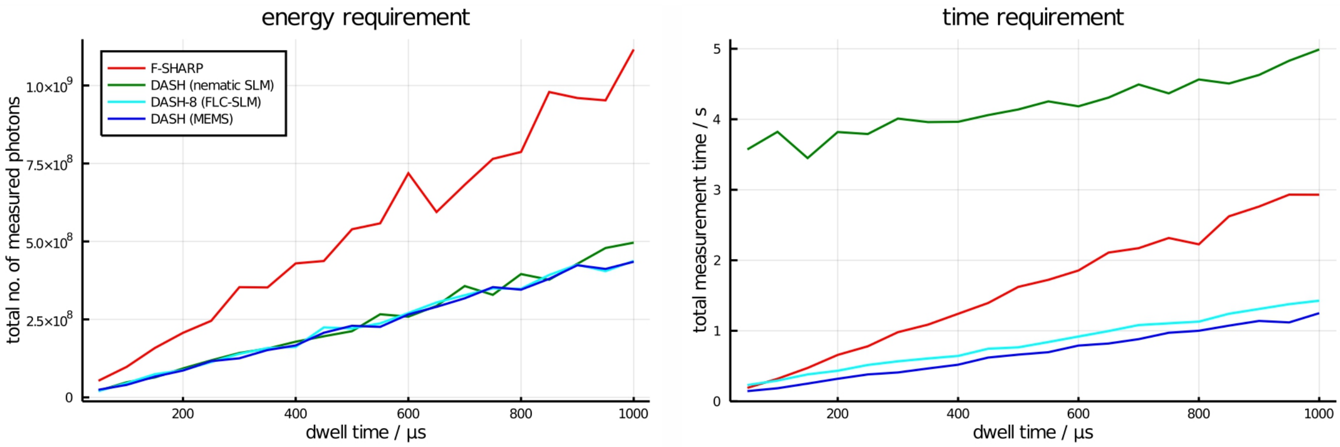


**Figure S1 | Photon budgets and correction time.** Number of photons required (left) and total measurement time (right) to establish a correction of 50% for increasing pixel dwell times.

The plots on the right show the total correction measurement times required for different pixel dwell times, that is the time the focus dwells on a particular position at the sample for signal acquisition. Naturally, the measurement times have a linear relation to the dwell times, albeit with different slopes and offsets. The slope for F-SHARP is about twice as high as for the DASH implementations, which means that DASH generally benefits if longer pixel dwell times are required. At a dwell time of 50 µs, the MEMS implementation of DASH performs approximately equally fast as F-SHARP. Analogously, the ferroelectric LC-SLM implementation is faster than F-SHARP for dwell times greater than 100 µs. Due to its comparably slow switching speed, the nematic SLM implementation of DASH is the slowest for the shown dwell time range. Nevertheless, the total correction times are in the range of a few seconds, which is likely to be sufficient for a number of different imaging experiments on live tissue [1].

The initial demonstrations of DASH shown in this work were made with a relatively slow nematic LC-SLM (Hamamatsu X10468-07) with a response time of 50 ms. In this case, nearly 10 minutes were required to optimize the DASH correction mask under the experimental conditions in Fig. 3 of the main text (10 iterations, 224 modes, and pixel dwell time 1 ms). This measurement time is, of course, much longer than the persistence times of most living tissues and therefore only useful for an initial demonstration of the technique. However, the SLM was then upgraded to a faster nematic LC-SLM (Meadowlark HSP1920) with a response time of 3 ms and the measurement time was dramatically reduced to just 45 seconds for the same experimental parameters compared to 22 seconds required for F-SHARP. Notably, DASH has the potential to further improve in speed by more than an order of magnitude if a fast MEMS mirror is used instead of the nematic LC-SLM (see Fig. S1). In this case, the required dwell time would become the rate limiting factor in most biological imaging experiments, and any remaining speed advantage of F-SHARP would be far outweighed by its requirement for higher photon numbers as emphasized in Fig. S1.

### II. EQUIVALENCE OF IMPACT AND F-SHARP

In this section, we provide a detailed derivation of the equivalence of IMPACT and F-SHARP.

The core idea of IMPACT is that many pixels are simultaneously modulated with increasing frequencies and interfered with a static reference wave behind the scatterer to produce oscillations in the TPEF signal. If the modulation frequencies are chosen appropriately, a single Fourier transform of the time-dependent signal yields the correction phases which are subtracted from the mask on the SLM after each iteration.

While the original implementation of IMPACT [2] alternately modulates only one half of the pixels and uses the static field of the others as reference, it is also possible to modulate all pixels simultaneously and to provide the reference wave externally. Although this is not necessarily more practical, we shall assume this case to facilitate the following considerations. We further assume a modulator featuring $N$ pixels, which are modulated by unique frequencies $\omega_n = N\,\omega_0 + n\,\omega_0$, where $n$ is the pixel index and $\omega_0$ a certain frequency that can be freely chosen. At an arbitrary point behind the scatterer, the fields of all pixels $u_n e^{j\omega_n t+\phi_n}$ interfere with each other as well as the static reference wave $re^{j\varphi}$. Assuming scalar fields, the intensity at this point can be expressed as:

$$I_t \propto \left(\sum_{n=0}^{N-1} u_n\, e^{j\omega_n t + j\phi_n} + re^{j\varphi}\right)\left(\sum_{m=0}^{N-1} u_m e^{-j\omega_m t - j\phi_m} + re^{-j\varphi}\right) = \\ = r^2 + A_t + B_t\, e^{-j\varphi} + B_t^*\, e^{+j\varphi}, \tag{1}$$

with

$$A_t = \sum_{n=0}^{N-1}\sum_{m=0}^{N-1} u_n\, u_m\, e^{j(n-m)\omega_0 t + j(\phi_n - \phi_m)} \tag{2}$$

$$B_t = r\sum_{n=0}^{N-1} u_n\, e^{j(n+N)\omega_0 t + j\phi_n}. \tag{3}$$

Note that the time $t$ is expressed as an index to emphasize its discrete character. The frequency spectrum of $I_t$ reveals that our particular choice of modulation frequencies $\omega_n$ ensures a clear separation of the spectrum of $B_t$, which contains the desired phases $\phi_n$, from the contributions of $B_t^*$ and the mixing terms of $A_t$. Therefore, the phases can be extracted by computing a discrete Fourier transform of the recorded time signal and considering only the relevant entries of the result. According to the Nyquist theorem, $4N$ sampling points are required to compute $2N$ phase values, half of which are finally used. This is the path followed in Ref. [2].

We note that taking $3N$ measurements is actually sufficient if one considers a phase stepping scheme, where $N$ measurements are taken for each one of three equidistantly chosen reference phases: $\varphi = [0, 2\pi/3, 4\pi/3]$. From Eq. 1 it becomes apparent that a value proportional to $B_t$ can be retrieved by multiplying the three time signals with the conjugates of the respective reference phase factors and averaging the result:

$$B_t \propto \frac{1}{3} \sum_{p=0}^{2} I_{t,p} \; e^{-j\varphi_p}. \tag{4}$$

Notably, the separation of desired phase terms in frequency space is not a requirement for the phase-stepping scheme, which is why we can choose the modulation frequencies to be $\omega_n = n\ \omega_0$. The phases $\phi_n$ are then contained in the phase part of the discrete Fourier Transform of $B$:

$$\phi_n = \mathrm{angle}\left(\mathrm{DFT}[B_t]\right). \tag{5}$$

On the other hand, in F-SHARP a weak external beam is scanned across a stronger, phase corrected static field. The stronger corrected field is assumed to form a more tightly confined focus behind the scatterer due to the nonlinearity of the two-photon signal, and thus takes the role of the "probe beam". Interferograms are then measured for different phases of the scanning beam and the amplitude and phase at each position of the scanning field are retrieved by the same phase stepping algorithm used in DASH. The final phase mask is retrieved by numerically propagating this measured field to the SLM plane and taking its phase.

For the following considerations we are assuming a rectangular pupil comprising $N = N_x \cdot N_y$ pixels on the SLM and an equally shaped grid of scanpoints in the focal plane. We further assume a typical sawtooth galvo motion with a fast x-axis and a slow y-axis, respectively. The galvo sweeps the weak beam across the focal plane. With no scatterer in place, the focused field takes the form of a translated delta function: $\delta(x - \Delta s_x \ \mathrm{mod}_{\mathrm{Nx}}[t], y - \Delta s_y / N_y \ t)$. Note that $t$ is a dimensionless index ranging from 0 to $N_x \cdot N_y - 1$. An optimal choice for the step sizes $\Delta s_{x,y}$ is governed by the Nyquist theorem: $\Delta s_{x,y} = \lambda/(2\mathrm{NA}_{\mathrm{x,y}})$, where $\mathrm{NA}_{\mathrm{x,y}}$ denote the numerical apertures along the x- and y-directions. For this case, the phase of the field in the Fourier plane, at the positions of the SLM pixels, is:

$$\phi_{m_x,m_y,t} = \frac{2\pi}{N_x}\ m_x \ \mathrm{mod}_{\mathrm{Nx}}[t] + \frac{2\pi}{N_y}\frac{m_y}{N_y}\ t, \tag{6}$$

where $m_x$ and $m_y$ are column and row indices of the SLM pixels. From this equation we see that the mirror fly-backs after the end of each line scan do not introduce any phase discontinuities. In these cases, $\mathrm{mod}_{\mathrm{Nx}}[t]$ changes from $N_x - 1$ to 0, which, on top of adding the desired increment $2\pi/N_x \cdot m_x$ merely subtracts multiple integers of $2\pi$ from the phases of every pixel. Therefore, we can neglect the modulo operator and rewrite Eq. 6 using a linear pixel index $n = m_y + m_x\ N_y$:

$$\phi_{n,t} = \frac{2\pi}{N} n\ t, \tag{7}$$

which allows us to identify unique modulation frequencies for each pixel: $\omega_n = \frac{2\pi}{N} n$. We can thus conclude that the scanning motion used in F-SHARP can be interpreted as a particular multidither scheme of IMPACT.

The performance of IMPACT was also compared with DASH experimentally and the behavior was similar to that of F-SHARP. In these experiments, IMPACT was implemented as described in Kong, et. al. [3] by modulating a randomly selected half of in total 256 SLM pixels at once. However, as discussed above, modified frequency assignments and phase stepping were used to reduce the required number of measurements.

TPEF images of a 4 µm fluorescent bead under a scattering mask are shown in Fig. S2 with no correction (a), DASH correction (b), and IMPACT correction (c) after ten iterations each, along with the measured correction masks (insets). Similarly to F-SHARP, the final enhancement using IMPACT was somewhat lower than for DASH, as illustrated by the intensity profiles in Fig. S2(d). This could be due to several factors, including the use of power splitting $f = 0.5$ which is implicitly made by modulating half of the pixels, or the choice of a pixel mode basis instead of a Fourier basis.

The signal enhancements during each measurement for the two algorithms are also shown in Fig. S2(e). Here, the IMPACT enhancement is consistently lower than that of DASH because only half of the SLM pixels are corrected during each measurement while the other half are modulated, while in DASH 80% of the power was in the corrected beam. Note that this does not affect the final correction like that shown in Fig. S2(d), where the optimal phase is applied to all of the SLM pixels.

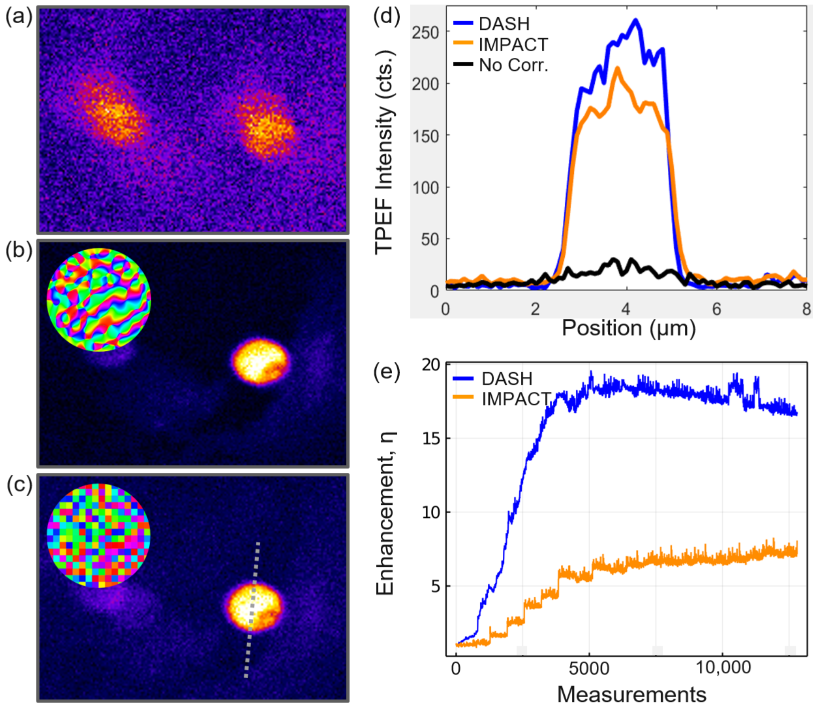


**Figure S2 | Experimental comparison of IMPACT and DASH**. (a)-(c) Images of a 4 µm fluorescent bead under scattering tape taken with no correction, with DASH correction, and with IMPACT correction, respectively. Correction masks for DASH and IMPACT are shown as insets. (d) Intensity profiles along the dashed line in part (c) for each image. (e) TPEF enhancement at each measurement throughout the algorithms.

### III. OPTIMAL VALUES OF THE POWER SPLITTING RATIO $f$

The factor $f$ is defined as the power fraction used for the modulated wave and thus it determines how the total power is shared between the reference and modulated waves. At the beginning of the algorithm, when both waves are equally scrambled by the scatterer, a value of $f = 0.5$ maximizes the power in the signal $B_t$ and $B_t^*$, which is proportional to $r\,|U|$, with $|U| = \sum_n |u_n|$. This can be deduced from Eq. 3.

However, in DASH, the reference wave becomes gradually better focused as the algorithm proceeds, and the convergence can be improved by choosing $f > 0.5$ to maintain an optimal power balance between reference and modulated waves. This was confirmed in the simulations shown in Fig. S3 (left) which visualize the influence of the power splitting ratio $f$ on the two-photon signal enhancements after one iteration of DASH and F-SHARP for a single fluorescent emitter in the focal plane assuming 4096 scattering and 256 correctable modes and 1000 photons/measurement as the initial signal level. Under these conditions, an optimal value of $f = 0.6$ was found for DASH. Conversely, F-SHARP obtains the largest two-photon signal enhancement with $f = 0.5$ after the first iteration, because at this point each mode has been tested once but no correction has yet been applied.

Importantly, Eq. 3 formulates the signal created at *one single point* behind the scatterer and not the actually recorded PMT data in a realistic application scenario, where the signals of all fluorescing points are accumulated. For an extended fluorescent sample, many speckles are created in the focal plane and the brightest point will eventually dominate and determine the position of the future focus [4]. In this context it is important that the constant reference wave determines the brightest speckle and not the continuously changing modulated wave, otherwise the algorithm is unlikely to converge onto a single stationary point. This is facilitated by choosing the reference wave to be more powerful than the modulated wave, i.e., $f < 0.5$ as shown in the right panel of Fig. S3 where the optimal values for $f$ lie below 0.5 for both F-SHARP and DASH. Here, the simulated correction was continued up to the 5th (F-SHARP) and 3rd (DASH) iterations, to obtain sufficiently large enhancement factors and initial signals of 10k (F-SHARP) and 1k (DASH) photons/measurement were assumed.

For an extended fluorescent sample, the optimal value of $f$ depends on a trade-off between minimizing $f$ to favor a focus determined by the constant reference wave and maintaining enough signal contrast between modulated and reference beams to make an accurate phase measurement. For this reason, the optimal value of $f$ is determined by the initial signal level as shown in Fig. S4(a), in which the enhancement after one simulated DASH iteration was investigated as a function of $f$ for increasing initial signals from a uniform 2D fluorescent layer. For the lowest signal

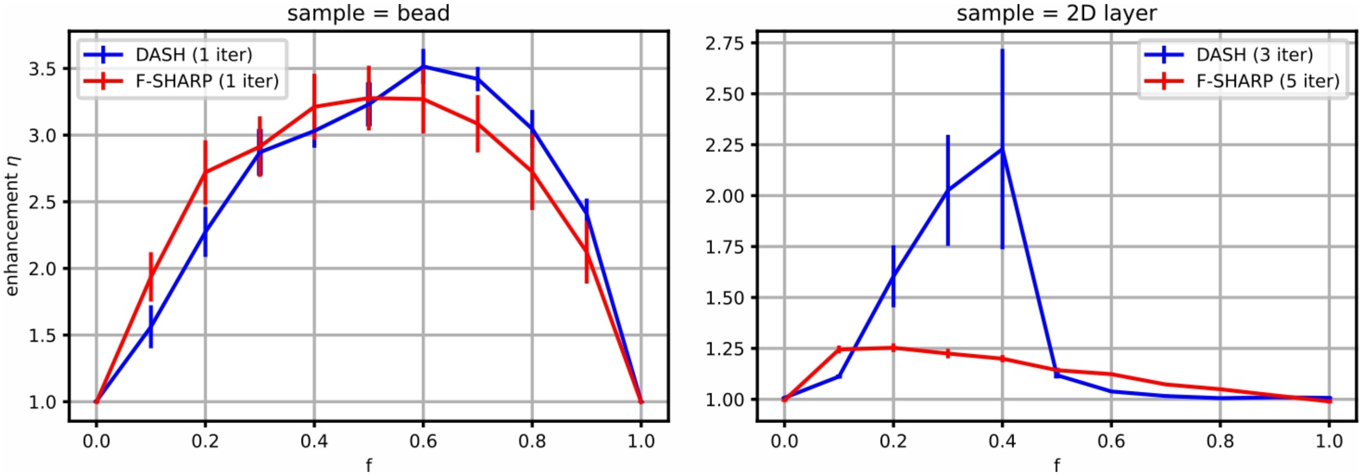


**Figure S3 | Influence of the power splitting ratio $f$ on the convergence behavior of F-SHARP and DASH**. Left: Optimizing the signal from a single fluorescent point; Right: Assuming a 2D fluorescent layer as sample; Each data point represents the average TPEF signal enhancement achieved after a certain number of iterations (stated in the figure legend). The mean values are based on 10 subsequent trials, the error bars mark their standard error.

(540 photons/measurement), the enhancement is optimized around $f = 0.3$, while for the highest signal (540k photons/measurement) the enhancement improves down to the smallest simulated ratio $f = 0.01$. This behavior is illustrated in Fig. S4(b)-(c) where the enhancement for each measurement in the low signal and high signal cases is plotted for three increasing values of $f$.

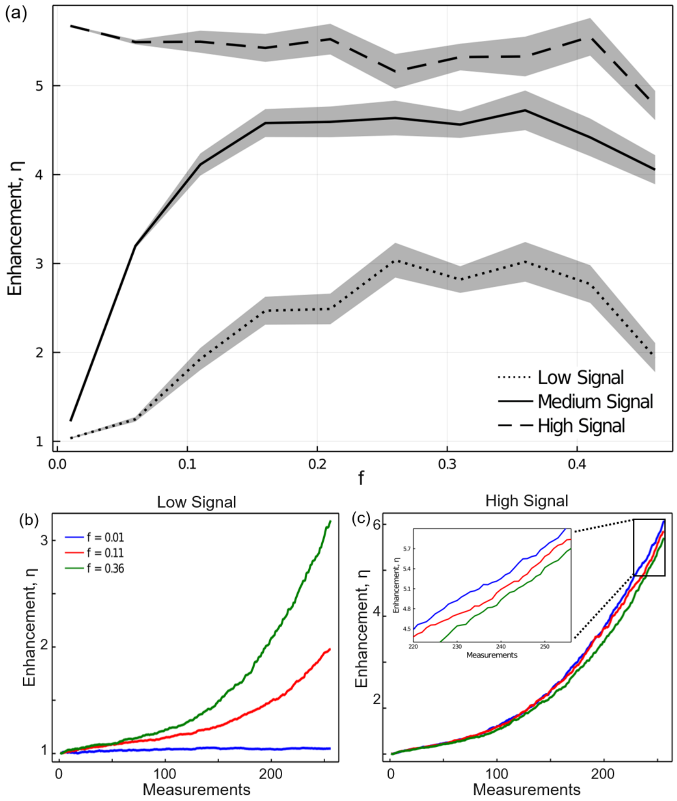


**Figure S4 | Optimizing $f$ for increasing signal levels**. (a) Mean signal enhancement after the first DASH iteration for low, medium, and high initial signal levels corresponding to 540, 2700, and 540k photons with standard error of the mean over 10 trials shown as ribbons. (b)-(c) Signal evolution for $f = 0.01$, $f = 0.11$, and $f = 0.36$ with low and high initial signals, respectively.

### IV. PHASE STEPPING ALGORITHM

A well established phase stepping interferometry algorithm was used to rapidly determine the phase $\phi_{i,n}$ and amplitude weighting $a_{i,n}$ for each mode $M_n$ from the 2-photon fluorescence signals $S_{i,n,p}$ for each phase $p$ using the equations shown below.

$$\phi_{i,n} = \text{angle}\left(\sum_{p=0}^{P-1} \sqrt{S_{i,n,p}}\; e^{-j\varphi_p}\right) \tag{8}$$

$$a_{i,n} = \frac{1}{r_{i,n}}\frac{1}{P}\text{abs}\left(\sum_{p=0}^{P-1} \sqrt{S_{i,n,p}}\; e^{-j\varphi_p}\right) \tag{9}$$

where $P$ is the total number of measured phases and $r_{i,n}$ is the amplitude of the reference wave at iteration $i$ and mode number $n$. Note that $r_{i,n}$ at our considered point behind the scatterer is usually unknown. This is unproblematic if the SLM is only updated after each full measurement iteration comprising all modes, such as in F-SHARP, because $r_{i,n}$ does not change during an iteration and only relative weightings between different mode amplitudes $a_{i,n}$ matter. Then, $r_{i,n}$ can for instance assumed to be 1 in Eq. 9. For DASH, however, $r_{i,n}$ continuously increases during an iteration, because the reference beam becomes increasingly better corrected. Therefore, it is helpful to consider at least an estimate of $r_{i,n}$, which we can obtain from the 2-photon fluorescence signals measured in the phase stepping routine. Since the square root of a two-photon signal is proportional to the excitation intensity expressed in Eq. 1, we conclude that

$$\sum_{p=0}^{P-1} \sqrt{S_{i,n,p}} \propto r_{i,n}^2 + A_n, \tag{10}$$

where $r_{i,n}^2$ and $A_n$ represent the excitation intensities of reference beam and modulated beam at the point behind the scatterer that we consider here, and by summing over the phases the cross terms $B$ average to zero. If both beams are equally attenuated by the scatterer, their intensity ratio at the very first measurement is $r_{i,n}^2/A_n = (1-f)/f$, which is about 2.3 for $f = 0.3$. As the algorithm proceeds, $r_{i,n}^2$ increasingly dominates even more over $A_n$, which is why we can neglect $A_n$ in Eq. 10 and calculate $r_{i,n}$ as:

$$r_{i,n} = \left(\sum_{p=0}^{P-1} \sqrt{S_{i,n,p}}\right)^{1/2}. \tag{11}$$

Adjusting $r_n$ after each measurement only modestly increases the convergence rate, so this effect was neglected in our experiments and a constant value $r_n = 1$ was assumed.

It is also worth noting that while theoretically P=3 phase steps are enough for the interferometric phase measurement, a measurement performed with this minimum number is highly susceptible to environmental factors like time dependent background signals. The accuracy of the interferometric measurement can be improved by increasing the number of phase steps, but at the cost of an overall increase in the measurement time. Here, this trade-off was optimized experimentally by performing DASH on a test sample with increasing numbers of phase steps and choosing the number (P=5) that gave the highest enhancement after 1000 total measurements. This choice was further confirmed by fitting the measured two-photon data to a cosine function and increasing the number of phase measurements until the sum shown in Eqn. 8 returned the same phase offset as directly fitting the data.

### V. GENETIC ALGORITHM

The GA used for the comparison in Fig. 1 of the main document was implemented following the procedure in Ref. [5]. In contrast to the other methods, the performance of the GA depends on several parameters, which have been manually optimized for the given boundary conditions in many subsequent simulation trials. Consistent with the nomenclature and variable definition in Ref. [5], the GA is initialized with a population comprising 50 random phase patterns. These initial patterns are tested and ranked according to their corresponding two-photon fluorescence signals. Then, the algorithm is executed for 300 iterations ("generations"). In each iteration 25 "offspring" patterns are generated, each derived from two different "parent" patterns of the former generation, by combining 50% of the pixels (randomly selected) of each parent. The parent selection is governed by a probability distribution which prefers well performing patterns yielding a high two-photon signal. The probability to choose pattern $i$ in the ranking is proportional to $(S_i - S_{50})^5$, where $S_i$ and $S_{50}$ are the two-photon signals obtained with pattern $i$ and the worst performing pattern in the ranking. A certain fraction of phase pixels is then randomly changed in each offspring pattern. This mutating fraction is continuously decreasing over the course of the algorithm by an exponential law: $(R_0 - R_{end})\exp(-g/\lambda) + R_{end}$, where we have chosen $R_0 = 0.02, R_{end} = 0.01, \lambda = 10$. $g$ denotes the index of the generation. Finally, one iteration is concluded by replacing the 25 worst-performing parent patterns by all of the offspring patterns and ranking the entire set, which represents the next generation.

### VI. ANIMALS AND TISSUE PREPARATION

All procedures involving animals were carried out in accordance with the Ethics Guidelines of Animal Care (Medical University of Innsbruck), as well as the European Communities Council Directive of 22 September 2010 on the protection of animals used for scientific purposes (2010/63/EU), and approved by the Austrian National Animal Experiment Ethics Committee of the Austrian Bundesministerium für Bildung, Wissenschaft und Forschung (permit number BMBWF-66.011/0148-V/3b/2019).

Mice expressing enhanced green fluorescent protein (GFP) under the promoter of Cx3cr1 (Cx3cr1$^{GFP}$ mice, The Jackson Laboratory: 005582) were used to visualize microglial cells in fixed brain slices. Animals were housed under specific pathogen-free (SPF) conditions at constant room temperature of 24°C on a 12h light/dark cycle with lights on from 07:00 to 19:00 and had ad libitum access to autoclaved pelleted food and water.

One heterozygous male Cx3cr1$^{GFP/+}$ mouse (20 weeks) was anesthetized with a mixture of ketamine (Ketasol®, 20 mg/ml) and xylazine (Xylasol®, 2 mg/ml) in 0.01M phosphate buffered saline (1X DPBS, 5µl/g body weight, i.p.). The mouse was transcardially perfused with 30 ml DPBS followed by 30 ml of ice-cold 4% paraformaldehyde (PFA), the brain was removed and postfixed in 4% PFA for two hours on ice. For brain slice preparation, the brain was further trimmed with a scalpel blade and glued onto the stage of a vibrating microtome (VT1000S, Leica Microsystems). Coronal slices (thickness 600 µm) containing the hippocampus were cut in PBS and subsequently stored in DPBS + 0.05% $NaN_3$. Slices were mounted onto microscope slides, embedded in Mowiol 4-88 and coverslipped. Slides were dried over night at room temperature and coverslips sealed using transparent nail polish.

### VII. LOW SIGNAL CONVERGENCE OF DASH

Even with an initial signal of 25 photons/measurement, the DASH algorithm converges to an enhancement of $\eta \sim 22$ after 50 measurement iterations as shown in the simulation results below.

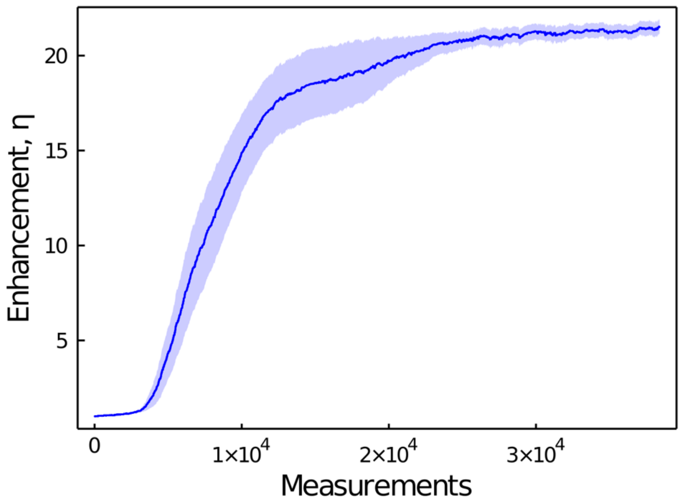


**Figure S5 | Convergence of DASH under low signal conditions**. Simulated DASH enhancement over 50 measurement iterations for an initial signal of 25 photons/measurement.
